# Co-Expression Network-based Analysis associated with potato initial resistance

**DOI:** 10.1101/496075

**Authors:** Lang Yan, Xianjun Lai, Yan Wu, Xuemei Tan, Haiyan Wang, Yizheng Zhang

**Affiliations:** College of Life Sciences, Sichuan University, Key Laboratory of Bio-resources and Eco-environment, Ministry of Education, Sichuan Key Laboratory of Molecular Biology and Biotechnology, Center for Functional Genomics and Bioinformatics, Chengdu 610065, Sichuan, China; Laboratory of Potato Functional Genome and Application, Xichang College, Liangshan, 615000, Sichuan, China

**Keywords:** potato, RNA-seq, gene expression patterns, co-expressing genes

## Abstract

RNA sequencing (RNA-seq) providing genome-wide expression datasets has been successfully used to study gene expression patterns and regulation mechanism among multiple samples. Gene co-expression networks (GCNs) studies within or across species showed that coordinated genes in expression patterns are often functionally related. For potatoes, a large amount of publicly available transcriptome datasets have been generated but an optimal GCN detecting expression patterns in different genotypes, tissues and environmental conditions, is lacking. We constructed a potato GCN using 16 published RNA-Seq datasets covering 11 cultivars from native habitat worldwide. The correlations of gene expression were assessed pair-wisely and biologically meaningful gene modules which are highly connected in GCN were identified. One of the primitively native-farmer-selected cultivars in the Andes, ssp.*Andigena*, had relative far distance in gene expression patterns with other modern varieties. GCN in further enriched 134 highly and specifically co-expressed genes in ssp.*Andigena* associated with potato disease and stress resistance, which underlying the dramatic shift in evolutionary pressures during potato artificial domestication. In total, the network was consisted of into 14 gene models that involves in a variety of plant processes, which sheds light on how gene modules organized intra- and inter-varieties in the context of evolutionary divergence and provides a basis of information resource for potato gene functional studies.

## Introduction

As the fourth highest food crop with global production over 350 million tons, tropical-originated potato has been cultivated worldwide nowadays with a wide variety of habitats ranging from cool highland zones to tropical lowlands Pino et al. (2007); Kikuchi et al. (2015); Hawkes et al. (1990, 1994). Although applied over relatively short timescales, artificial selection has dramatically altered the tuber sizes and shapes, color in skins and fleshes, providing a wide range of germplasm resources accessible for research and breeding purposes Machida-Hirano (2015). Along with the publishing of potato reference genome, represented by the doubled monoploid *Solanum tuberosum* group *Phureja* clone Consortium et al. (2011), researchers have insight into the genetic performances in potatoes, from basic gene structure and biological function to their expression profiles. While advances over the past years have generated a growing number of gene expression datasets which provide unbiased snapshots of gene expression dynamics among worldwide potato cultivars, it remains a key challenge to deduce the underlying gene regulatory circuits in the process of the potato genetic diversification.

To address this, ever more gene expression datasets can be merged and integrated to develop a system-wide research Hughes et al. (2000); Wu et al. (2002); Childs et al. (2011). Gene co-expression networks (GCNs) has emerged as a tool that incorporated large-scale gene expression analyses, which describing the relatedness between gene expression patterns in a pairwise fashion Schmid et al. (2005); Lee et al. (2009); Wilkins et al. (2009). Utilizing Pearson correlation coefficient (PCC) to measure expression correlation between gene pairs from individual experiment, GCNs assembled via the Weighted Gene Coexpression Network Analysis (WGCNA) method could cluster gene pairs with PCC larger than a chosen threshold value into modules Langfelder and Horvath (2008). According to a ‘guilt-by-association’ paradigm, genes with associated biological functions often have similar expression patterns, the genes (nodes) and connections (edges) in the module could represent co-expression dynamics in different gene expression datasets. Although several GCN studies have been done, such as COB Schaefer et al. (2014), CORNET De Bodt et al. (2012) in maize and SCoPNet Bassel et al. (2011) in Arabidopsis, most of them integrated the microarray expression datasets of cultivars worldwide. Given that RNA-Seq has become the favored technique for detecting genome-wide expression patterns, the increasing number of RNA-Seq in potatoes implies an RNA-Seq based potato GCN protocol would be valuable to the scientific community Crookshanks et al. (2001); Ronning et al. (2003); Flinn et al. (2005); Rensink et al. (2005); Li et al. (2006); Wang et al. (2009).

Here, we integrated 16 published RNA-Seq datasets from 11 distinct potato cultivars worldwide to identify divergences of general gene expression patterns intra- and inter-varieties. Based on the correlations of gene expression patterns, we constructed a potato WGCNA-based gene co-expression network, containing 12,346 genes and assembling into 14 distinct gene modules with an average module size of 881 genes. This potato GCN enriched genes highly expressed and correlated to certain cultivars or tissues into modules, which highlighting functions in potato disease and stress responses, photomorphogenesis and tuber dormant release development .etc. Importantly, these identified modules could further used to analyze the transcription factors which acting as regulators controlling gene expression patterns. The potato GCN study could provide us the valuable resources with potato gene expression rules, help design proper crossing schemes in potato breeding across different geographical areas and be useful for potato systems biology analysis.

## Results

### Overview of RNA-Seq datasets

Publicly available potato gene expression data were downloaded from NCBI bio-project database, a collection of diverse biological data across multiple potato varieties and developmental tissues Gong et al. (2015); Campbell et al. (2014); Petek et al. (2014); Liu et al. (2015b,a); Lin et al. (2015). Only data including single- or pair-end RNA-Seq runs in the last five years were considered and there were totally 16 datasets with above 3 replicates remained, which monitored potato transcriptome from various cultivars and tissues under normal growth conditions (Table 1). In particular, four datasets in leaves represented distinct varieties as ‘Deisree’, ‘Sarpo Mira’, ‘Igor’ and ‘SW92-1015’, and another four were tubers in dormant and sprouting status of var.’Longshu3’ and var.’Russet Burbank’. The classic American commercial variety ‘Atlantic’, local Chinese varieties var.’Xindaping’ and var.’Heimeiren’ and primitive native-farmer-selected cultivars ssp.*Andigena* were also included. It is noteworthy that Andigena potatoes represents primitive cultivar group with a distinct gene pool compared to other modern potatoes, which used to identify widespread alteration of gene expression patterns during artificial domestication.

**Table 1.**
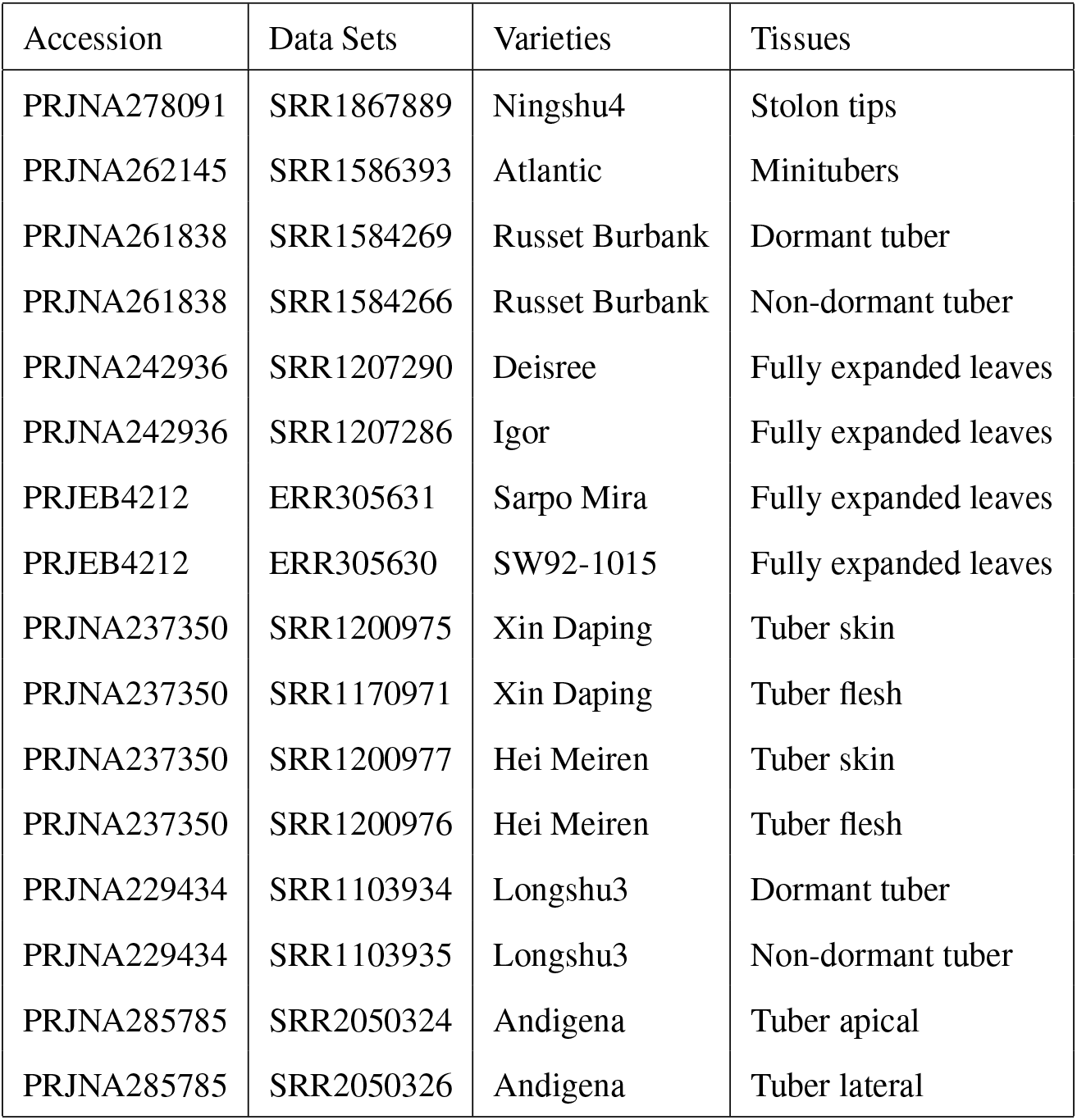
Potato gene expression datasets used in this study.

### Differential gene expression patterns between primitive and modern cultivars

The collected raw data files (.sra) were downloaded, mapped against to the *S. tuberosum* group *Phureja* DM v4.03 reference genome Trapnell et al. (2012); Consortium et al. (2011), and processed into gene expression matrix represented by Fragments Per Kilobase of transcript per Million mapped reads (FPKM)-normalized gene expression values. The genes with average FPKM values < 3.0 were filter out, resulting in a gene expression matrix involving in 12,346 genes (rows) and 16 runs (columns). After classifying gene expression values into negligible (FPKM < 0.5), extremely low (0.5 ⩽ FPKM < 3), low (3 ⩽ FPKM < 50), moderate (50 ⩽ FPKM < 100) and high (FPKM ⩾100), we found similar expression patterns in most modern cultivars, in which relatively less genes had negligible expression (191 genes in average) while a large number of genes had expression ranges from 3 to 50 FPKM (5,287 genes in average) (Figure 1). However, the primitive cultivar ssp.*Andigena* had distinct gene expression patterns in comparison with modern cultivars, even the comparisons were only involved in tubers. Nearly 3,000 genes in Andigena potatoes were tend to be silent with expression value < 0.5 FPKM and other 4,584 genes expressed extremely lowly which were in the range between 0.5 and 3 FPKM. The general expression pattern in primitive ssp.*Andigena* were leveled off without the expression peak in a certain region.

**Figure 1.**
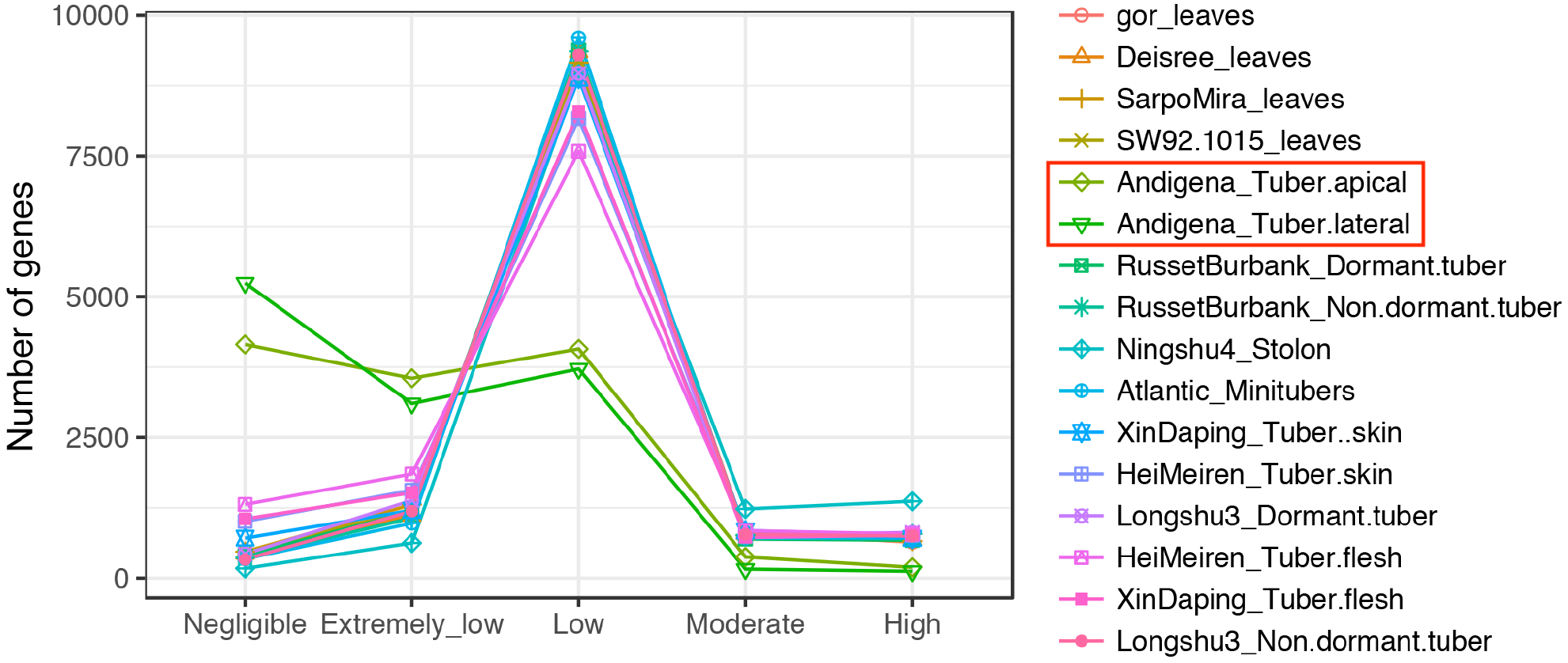
Gene expression patterns in different datasets. The genes with FPKM values were classified into negligible (FPKM < 0.5), extremely low (0.5 ⩽ FPKM < 3), low (3 ⩽ FPKM < 50), moderate (50 ⩽ FPKM < 100) and high expressed (FPKM ⩾100).

To identify the divergence of gene expression patterns in further between primitive and modern cultivars, the pair-wisely correlations of gene expression were calculated between datasets according to the spearman correlation coefficient (Figure 2, Supplementary Figure S1). The leaf tissues of 4 varieties (‘Deisree’, ‘Igor’, ‘SarpoMira’ and ‘SW92’) had the highest correlation coefficient between each pair ((r >0.85, p-values <0.05) compared to the rest tuber groups, indicating the higher correlations of gene expression patterns between tissues than that between cultivars (Table 2). Among tubers in different developing stage, gene expression had significant higher correlation coefficients within cultivars than that across cultivars, such as the dormant and non-dormant tubers of var.’RussetBurbank’ (r=0.88, p-value < 0.05) and that of var.’Longshu3’ (r=0.85, p-value <0.05). Some cultivars were close-related in breeding process or geographical area, resulting in the significantly correlated gene expression across cultivars, such as tuber skins between var.’Heimeiren’ and var.’Xindaping’ (r=0.89, p-value <0.05), tubers between var.’Atlantic’ and var.’RussetBurbank’ (r=0.83, p-value <0.05), tubers fleshes in var.’Xindaping’ and tubers in var.’Longshu3’ (r=0.86, p-value <0.05). Nevertheless, since the ssp.*Andigina* had no any significant correlations with other modern cultivars in expression levels (r < 0.65, p-value < 0.05), relationships between primitive and domesticated potatoes should be drifting apart.

**Figure 2.**
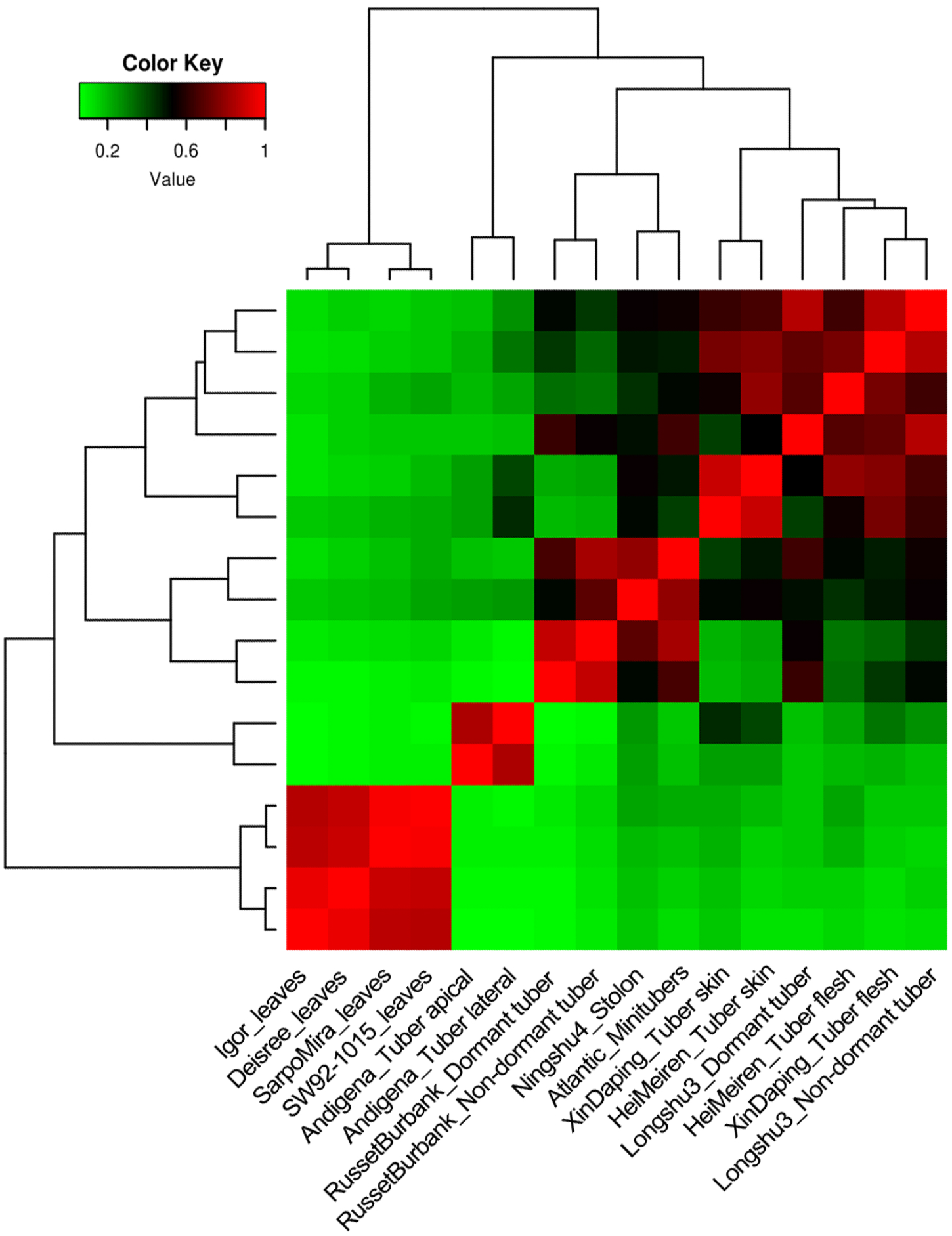
Cluster heat maps of log2-transformed FPKM values using the Spearman correlation coefficients among 16 datasets.

**Table 2.**
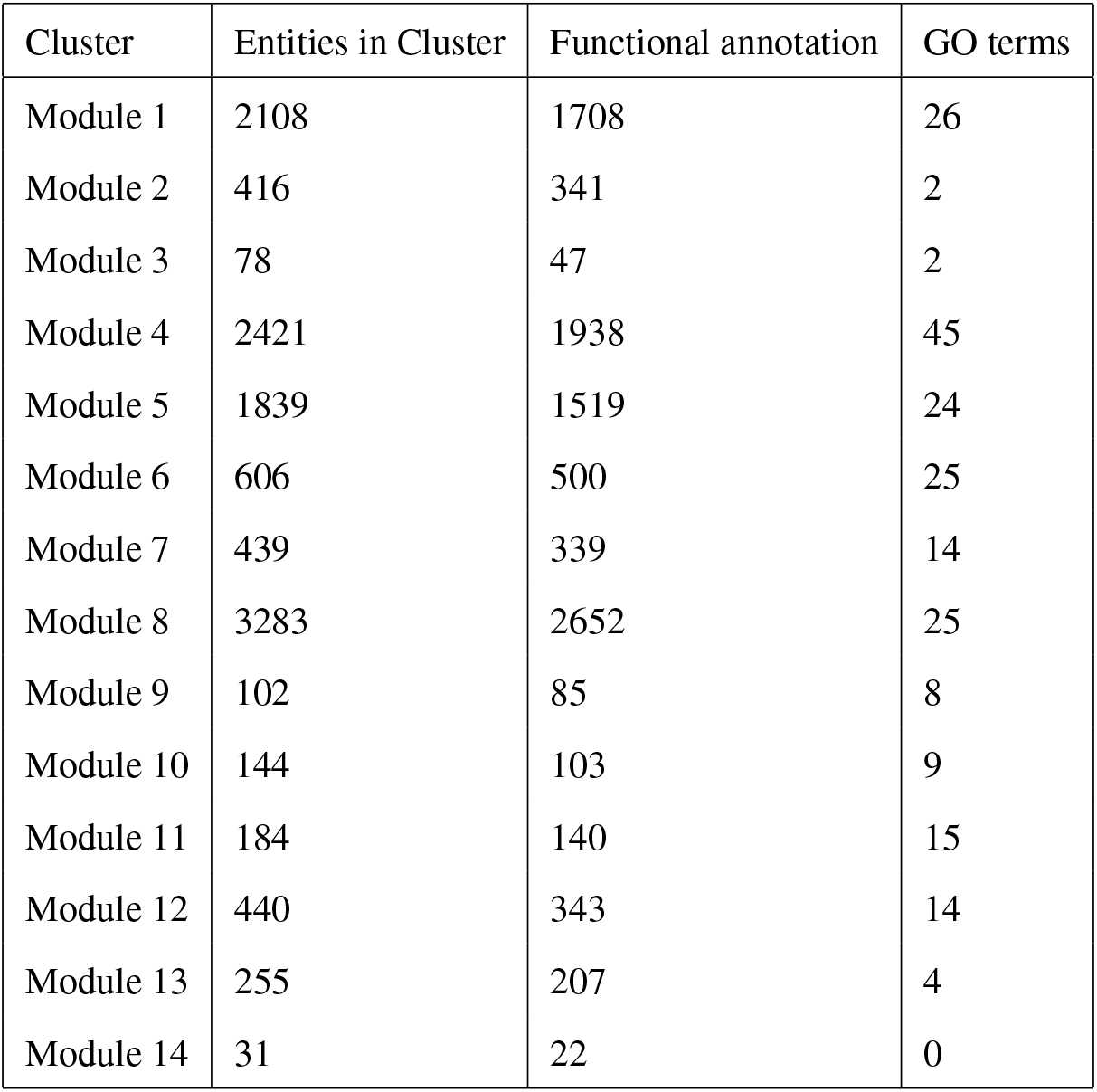
Summary of the number of entities and enriched GO terms in each validated module.

| Cluster | Entities in Cluster | Functional annotation | GO terms |
| --- | --- | --- | --- |
| Module 1 | 2108 | 1708 | 26 |
| Module 2 | 416 | 341 | 2 |
| Module 3 | 78 | 47 | 2 |
| Module 4 | 2421 | 1938 | 45 |
| Module 5 | 1839 | 1519 | 24 |
| Module 6 | 606 | 500 | 25 |
| Module 7 | 439 | 339 | 14 |
| Module 8 | 3283 | 2652 | 25 |
| Module 9 | 102 | 85 | 8 |
| Module 10 | 144 | 103 | 9 |
| Module 11 | 184 | 140 | 15 |
| Module 12 | 440 | 343 | 14 |
| Module 13 | 255 | 207 | 4 |
| Module 14 | 31 | 22 | 0 |

### Construction of gene co-expression modules

Based on correlations of gene expressions pair-wisely, GCNs were constructed using the weighted gene co-expression network analysis method (WGCNA), which is a systems biology approach aimed at understanding the gene expression patterns in networks instead of individual genes.Langfelder and Horvath (2008). The GCNs were consisted of 12,346 genes, involving in 14 distinct gene modules that contained 31-3283 genes with an average module size of 881 genes (Figure 3, Supplementary Figure S2). Each module represented genes with highly correlated expression profiles, either in a single or a few related datasets. The majority (80.5%) of genes assigned to co-expression modules have functional annotation. Thus, for genes with unknown functions, GCNs supplied potato gene annotation informations which placed unknown genes in a functional context and inferred their role according to their interaction with known meaningful genes. Functional enrichment analysis was performed on each module using total genes as population. A total of 213 GO terms were significantly enriched (Table 2). Since GO terms indicated the gene functional types in a certain module, it also contributed to understand the metabolic pathways which unknown genes involved in.

**Figure 3.**
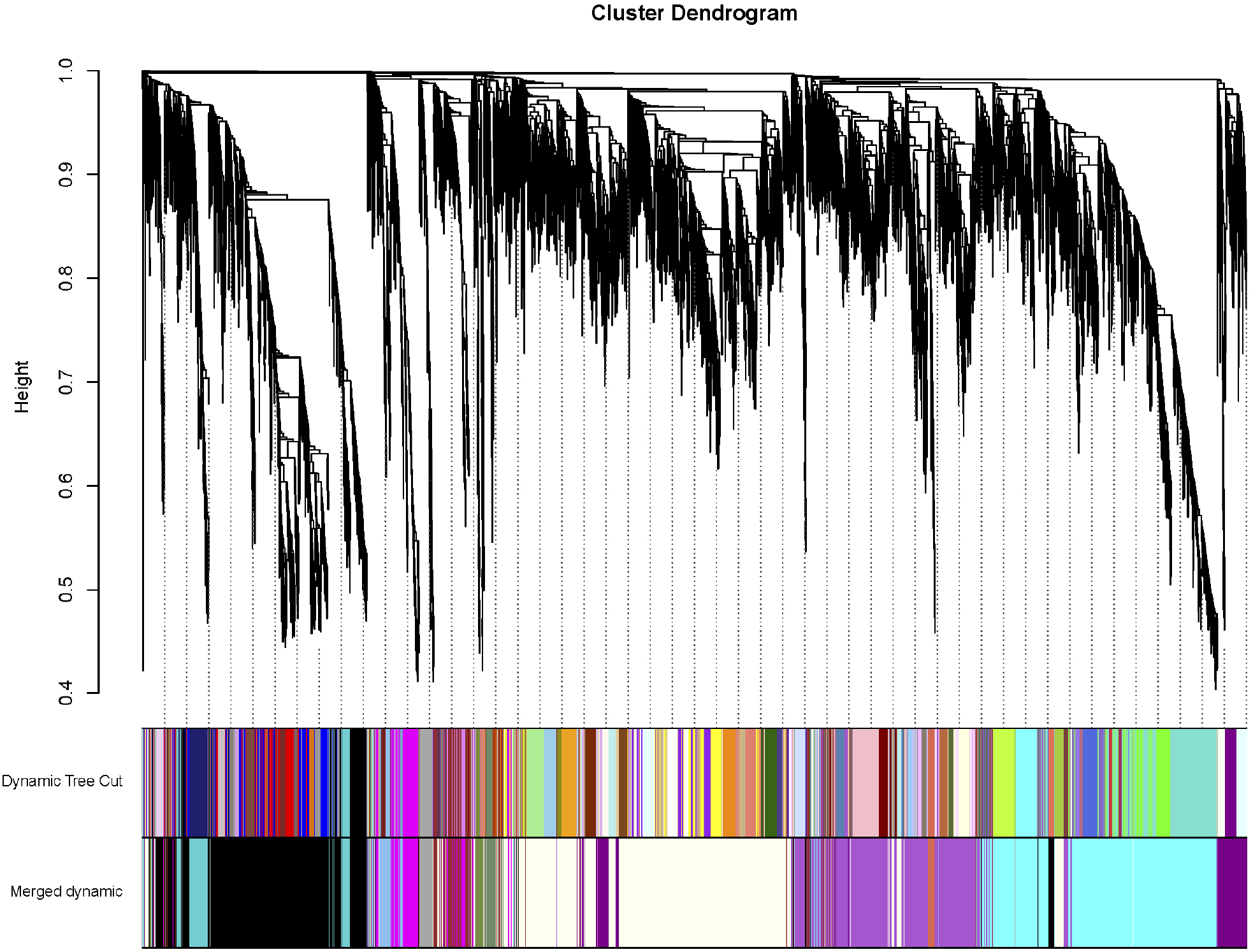
Hierarchical cluster tree showing co-expression modules identified by WGCNA. Each leaf in the tree represents one gene. The major tree branches constitute 14 modules are labeled by different colors.

Module eigengenes (MEs) for each module are the first principal component of a given module and can be considered as a representative of the module’s gene expression profileHollender et al. (2014). The MEs of 14 modules were calculated to check genes with similar expression patterns between different datasets, which showed in the heat map (Figure 4, Supplementary Figure S3). The dark red at the row-column intersection indicates the tissue specifically and highly expressed genes in the certain module. The result showed that gene expression patterns across different modules varied a lot. Six out of 14 modules are comprised of genes highly expressed in a single dataset (r >0.8, P < 10^−3^), which could be regarded as genes specifically correlated with a certain tissue/cultivar. Figure 5 displayed these tissue-specific gene expression modules which were mainly distributed on apical and lateral tubers of ssp.*Andigena* (Module #12 and #11), stolon of var.’Ningshu4’ (Module #4), non-dormant tuber of var.’RussetBurbank’ (Module #3) and tuber skin of var.’Heimeiren’ and ‘Xindaping’ (Module #9 and #10). In addition, some modules contained genes that were co-expressed highly in multiple datasets, implying that these genes could participate in particular regulations in multi-tissues/varieites. For example, Module #1 and #2 were both co-expressed highly in leaves and enriched in GO terms of cytochrome b6f complex (GO:0009512) and chloroplast part (GO:0044434), indicating the genes in these two modules were related to leaf development and morphogenesis. Module #7 and #8 were co-expressed highly in all of the datasets except in ssp.*Andigina*, corresponding with the hypothesis that some genes under selections during the domestication process had changed there general gene expression patterns.

**Figure 4.**
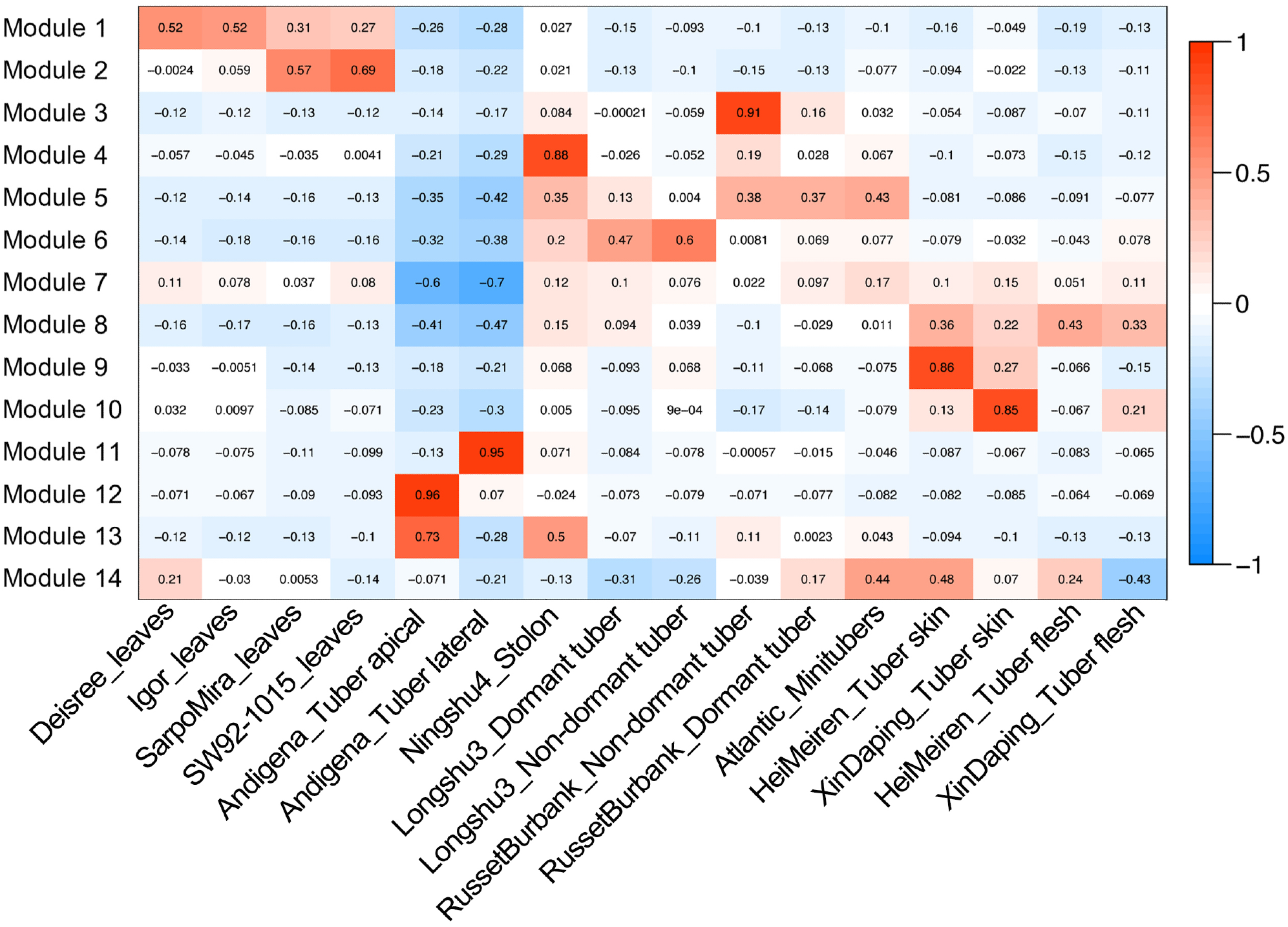
Heatmap representing the strength and significance of correlations between module eigengenes and different cultivars/tissues. Each row corresponds to a module. The number of genes in each module is indicated on the left. Each column corresponds to a specific tissue. The color of each cell at the row-column intersection indicates the MEs of the module in tissues.

**Figure 5.**
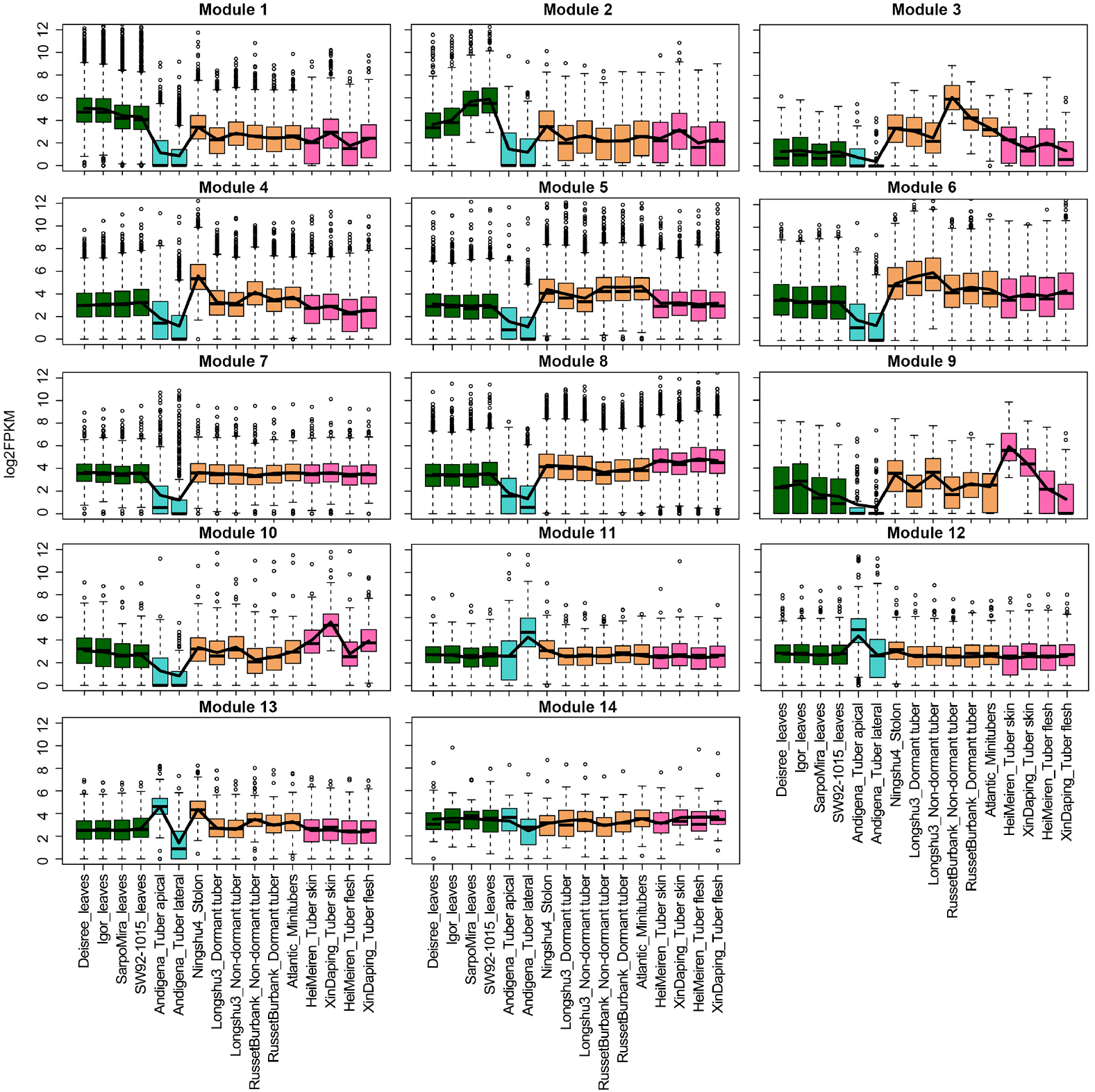
Boxplots of co-expression patterns of 16 datasets in the 14 modules. Gene expression values were processed using WGCNA to identify modules of highly correlated genes.

### Gene modules revealing certain developmental processes

The standardized MEs above were represented by the principal gene expression profile in modules, some of which showed the relatively high and specific expression level in certain tissues/cultivars. We then concerned about the biological processes of these tissue-specifically expressed genes participated in. Our GCNs identified four particularly relevant modules. Module #1 and #2 were enriched with genes involved in leaf-specific expression in multiple cultivars, and contributes to the process of leaf morphogenesis and photosynthesis, including chlorophyll biosynthetic and electron transport chain in cytochrome b6/f for transferring electrons within the cyclic electron transport pathway of photosynthesis activity. A number of genes for the development of photosystem were only included in these two modules: magnesium protoporphyrin IX methyltransferase (PGSC0003DMG400014243), light harvesting (PGSC0003DMG400020492) and G-protein coupled photoreceptor activity (PGSC0003DMG400010034), etc.

We firstly identified the expression pattern of genes enriched in Module 1 and 9, which had the tissue-specific expression in leaves of multiple varieties. Pair-wisely comparison of the gene expression difference and specificity were conducted in leaves among var.’Deisree’, var.’Igor’ and var.’Mira’ (var.’SW92’ was excluded due to lack of background information). The three varieties had 32, 33 and 77 specifically expressed genes (SEGs) respectively, which might be related to their varietal specificity like cultivated lineages and local habits etc. (Figure 6A). Among all the expressed genes, there were 63 differentially expressed genes (DEGs) between ‘Deisree’ and ‘Mira’, 83 between ‘Deisree’ and ‘Igor’, and 26 between ‘Igor’ and ‘Mira’. Moreover, there were some co-DEGs identified between two groups (Figure 6C,D), such as PGSC0003DMG400021142, the DGE in group ‘Deisree’ and ‘Mira’ (460.32 vs 2.22 FPKM) as well as the DGE between ‘Igor’ and ‘Mira’ (258.79 vs 2.22 FPKM). Importantly, two genes differentially expressed among all three cultivars, like PGSC0003DMG400011751, the expression value were 1881.08, 111.74 and 6.65 in ‘Deisree’, ‘Igor’ and ‘Mira’ respectively, indicating that the significant expressing volatility among different varieties. As expected, majority of the enriched genes in Module 1 and 9 of the co-expression network were DEGs and concentrated on photosynthesis process, including chlorophyll biosynthetic and electron transport chain in cytochrome b6/f for transferring electrons within the cyclic electron transport pathway of photosynthesis activity. The main functions of these genes encoded were magnesium protoporphyrin IX methyltransferase, light harvesting and G-protein coupled photoreceptor activity, etc.

**Figure 6.**
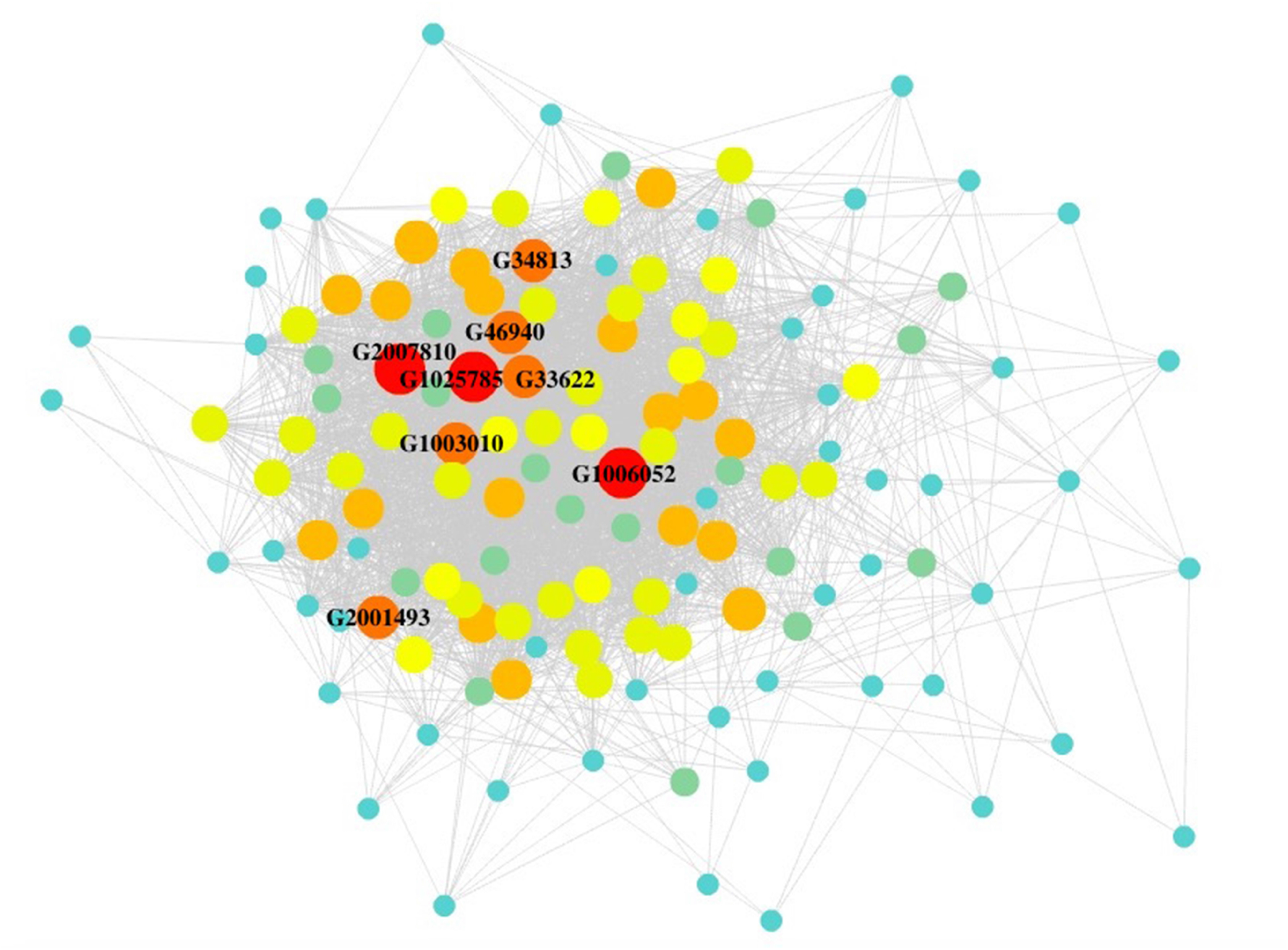
The correlation network 134 genes visualized by Cytoscape. Hub genes are indicated by larger circles shown in the network.

Compared to leaf morphogenesis, gene expression and regulation of tuber sprouting, which is a major yield-determining trait, attracted more attention of researchers. Two modules involved in tuber dormant releasing process were identified, providing us the principal regulatory candidates involved in the tuber sprouting development. Genes in Module #3 specifically and highly expressed in sprouting tubers of var.’Russet Burbank’, which predominantly encoded proteins such as auxin-induced protein, ethylene response factor, MADS-box protein, SAUR family protein, wound induced protein, etc. Seventy-three genes in Module #6 was specifically highly expressed in sprouting tubers of the Chinese local cultivar ‘Longshu3’, and showed functional specialization for tuber-specific and sucrose-responsive element binding factor, proteinase inhibitor, MYB domain class transcription factor, heat shock protein, etc.

Among the RNA-seqs in tubers, we compared the gene expression patterns of dormant and non-dormant tubers in both var.’Longshu3’ (‘LD’ and ‘LN’ for short) and var.’Russet Burbank’ (‘RD’ and ‘RN’), in which the genes related to dormant releasing process had enriched in Module 3 and 6 of the co-expression networks. Apart from the 11,594 genes expressed in all four tubers, there were more genes specifically expressed in sprouting than that in dormant tissues, demonstrating the increasing of the gene expression activities in the dormancy release process. Then, the comparison of gene differential expression pair-wisely showed 121 (LDvsLN), 68 (RDvsRN), 31 (LDvsRD), and 69 (LNvsRN) genes had significantly differential expression, respectively. It is noteworthy that there were totally 19 genes had strong variety-differences in expression levels since they had co-differential expression. Moreover, genes in module 3 and 6, which were specifically and highly expressed in sprouting tubers of var.’Russet Burbank’ and var.’Longshu3’ respectively, provided us the principal regulatory candidates involved in the tuber dormant release development. Although these two modules were all consisted of tuber-specific genes, their gene functions and metabolic process enriched had differences between varieties. Module 3 was predominantly enriched in proteins such as auxin-induced protein, ethylene response factor, MADS-box protein, SAUR family protein, wound induced protein, etc. while Module 6 was enriched in tuber-specific and sucrose-responsive element binding factor, proteinase inhibitor, MYB domain class transcription factor, heat shock protein, demonstrating the differential expression patterns in two varieties with distinct backgrounds.

### Primitive cultivar-specific module involves in initial resistance in ssp.*Andigena*

GCNs can be visualized as network maps using Cytoscape, in which the nodes represent genes and the connecting lines (edges) between genes represent gene correlationsShannon et al. (2003). Here we present an particular interest case of genes in a primitive cultivar-specific module (Module 11) that co-expressed specifically in lateral of ssp.*Andigena* (Figure 4). This network consisted of 134 correlative genes in which 3 hub genes had connections >80 and another 5 hub genes had connections >70 (Supplementary Table S1). Strikingly, 39 out of all hub genes (29%) were annotated as unknown functions and nearly half (10 out of 24) were functional unknown when only considering of the complicated regulatory genes with connections >60. These highly and specifically expressed genes with unknown functions in ssp.*Andigena*, which also existed in the primitive ssp.*Phureja* clone (the reference genome), might have correlations with the primitive cultivated potatoes under purifying selection during domestication.

The hub gene with the highest edge number (83 edges) is PGSC0003DMG401006052, a secA-type chloroplast protein transport factor, which is one of the important components in the protein translocation sec pathway in chloroplast. Other highly connected hub genes encoded guanine nucleotide regulatory protein, elicitor inducible LRR receptor, protein phosphatase 2c, senescence-associated protein, etc. Of particular importance is that lots of functional genes highly expressed in ssp.*Andigena* tended to be associated with potato disease and stress resistance. Previous studies have identified such genes in some model plants. For example, genes encoding the elicitor inducible LRR protein (EILP) was activated by treatment of salicylic acid or inoculation of *Pseudomonas syringae*, and the product of EILP amplified the sensitivity to disease stress and involved in non-host disease resistance in tobaccoTakemoto et al. (2000); the enhanced expression of senescence-associated genes (SAGs) could improve the ability in response to diseases caused by fungi, bacteria, and viruses through triggering the hypersensitive response (HR) or eliciting necrotic symptoms induced by virulent fungi and bacteria during infections, which had been observed in both the model plant *Arabidopsis thaliana* and a commercially important grapevine cultivarEspinoza et al. (2007); SAG29 in *Arabidopsis* could help the organism survive in the high salinity and other osmotic stress conditions through regulating the cell viability which may serve as a molecular link that integrates environmental stress responses into senescing processSeo et al. (2011). In addition, some functional genes with less connections in this network also showed important functions contributing to the potato growth and signal transduction processes such as gene encoding Auxin:hydrogen symporter, Auxin-induced protein X10A and F-box protein as an auxin receptor.

## Discussion

Gene expression data have expanded the availability of genetic resources, which not only had been used as the supplement for gene functional annotation in a genomic scale, but also help to detect the individual gene expression patterns among tissues, varieties, or environmental conditions. In this study, we combined the gene expression correlation analyses and gene co-expression network to identify the correlations and divergences of gene expression, which may imply common functions or regulatory pathways among multiple potato cultivars. Making use of the integrated public RNA-Seq datasets could contribute to the identification of highly correlated genes in multiple parallel datasets. It also provides us an opportunity to study the gene regulatory mechanisms related to some important biological processes in systematic level, which is unattainable in single or few transcriptomes. The GCN established in this study enriched the highly correlated and expressed genes in the processes of photomorphogenesis and tuber dormant release development as well as stress resistance related genes specifically and highly expressed in primitive ssp.*Andigena*. The gene expression divergence inter-varieties breeding from different native habitats worldwide had never been reported before.

Gene expression patterns analyses between different cultivars in this study provide us some clues in the future breeding. Firstly, gene expression correlation in parallel tissues across cultivars could be associate with their relationships during breeding process. As shown in the Table 2, the Chinese local cultivars, ‘Ningshu4’ and the American traditional commercial variety ‘Atlantic’, which were bred from different geographical area, had high correlation coefficient in their gene expression patterns, implying their close relationship in breeding. Var.’Atlantic’ should be the exotic breeding accessions in the breeding of ‘Ningshu4’. Secondly, this study supported the previous hypothesis that long-term storage of potatoes is largely associated with the genetic control in potato itself instead of external environmental factors like soil and weather during potato growthSuttle (2007). The GCNs identified two gene modules, both of which had enriched genes specifically and highly expressed in sprouting tubers of var.’Longshu3’ and var.’Russet Burbank’, respectively. The var.’Longshu3’ and var.’Russet Burbank’ were both self-stable with relatively long dormant period, belonging to mid-late maturity varieties. However, some dormancy-controlled genes and their regulated pathways during the dormancy release process had different expression patterns in two varieties. For example, the genes encoding pectinesterase and xyloglucan endotransglucosylase, which should be the indicators of metabolic shifting into cell wall loosening, were found to be up-regulated expression only in non-dormant tubers of var.’Russet Burbank’. As reported before, the high expression of xyloglucan endotransglucosylase linked with protein increase that alters turgor pressure or cell wall pH valueresulting in the cell expansion Van Sandt et al. (2007); Senning et al. (2010); De Vos et al. (2012); Wolf et al. (2012). The pectinesterase has also been shown to be associated with increase in cell wall H^+^ Wolf et al. (2012). Oppositely, non-specific lipid-transfer proteins and lipid binding proteins encoded genes were significantly highly-expressed and enriched in the dormant tubers of var.’Longshu3’ rather than var.’Russet Burbank’. As most of the genes required for lipid degradation were expressed before bud emergence, which lead to energy release and supply of carbon resources for the production of sucrose to bud growth, the related genes in var.’Longshu3’ might positively participate in lipid metabolism during the process of dormancy releasePalta et al. (1993); Liu et al. (2015b).

Moreover, our results supported the conclusion that long-term storage of potatoes is largely associated with the genetic control in the variety itself except for the external environmental factors like soil and weather during potato growthSuttle (2007). The var.’Longshu3’ and var.’Russet Burbank’ were both shelf-stable with relatively long dormant period, belonging to mid-late maturity varieties. The pair-wisely comparison of LD-LN and RD-RN demonstrated that although DEGs appeared during the dormancy release process, the dormancy-controlled genes and their regulated pathways between two varieties were not completely identical.

Lastly but most significantly, the specifically- and highly-expressed gene modules and their regulatory network in ssp.*Andigena* are valuable resources to identify the genetic locis being targets of artificial selection during potato domestication. Potatoes are native to the Andes of South America, where represent at best genetic diversity among potato germplasmSpooner et al. (2005a). Andigena *(Solanum tuberosum L*. subsp.*andigena Hawkes*) is the most primitive cultivated potato in the Andean highlands and is a likely ancestor of the worldwide grown modern potatoes *S.tuberosum* Hawkes et al. (1990); Hawkes (1956); Sukhotu and Hosaka (2006). Andigena potatoes hold genotypes of wild species and gradually form the abundant modern cultivated populations in ever-changing environment selected by nature and human beings, thus it is important in germplasm resources as a primary gene pool for improving worldwide grown potatoes in breedingSukhotu and Hosaka (2006). Along with the domestication and long-distance dispersal of seed tubers spreading from South America to Europe, Africa, Asia, and diffusing world widely, ssp.*Andigena* had shown multiple differences with the modern cultivars morphologically and physiologically, which are also reflected on the gene expression patternsGrun (1990); Spooner et al. (2005b); Ames and Spooner (2008). In our study, the expression of multiple gene in ssp.*Andigena* had decreased which is silenced or expressed extremely lowly. Thus, Andigena had no significant correlations with modern varieties in gene expression level, demonstrating the massive variation in transcriptional levels between primitive and modern cultivars. Moreover, Andigena remained valuable traits in resistance and nutritional contents which had lost in modern potato cultivars, not only conferring resistance to late blight, potato virus X, potato virus Y, nematodes, tuber moth, etc., but also acting as as a source of antioxidant phytochemicals and mineral micronutrients like carbohydrates, vitamin C, phenolic, carotenoid etc. Andre et al. (2007). For this reason, Andigena had its wide variability in tuber shape, flavor and cooking quality, which differ from the modern domesticated cultivarsAndre et al. (2007). As genetic improvement in modern potato breeding is facing increasingly narrow genetic basis and the decline of the genetic diversityChimote et al. (2004); Collares et al. (2004); Fernie et al. (2006), the introduction of primitive cultivars like Andigena, a very important gene pool, could provide the luxuriant and distinct gene resources for breeding and agricultural production. Although the multiple especially-correlated genes identified and enriched in Andigena showed a considerable variability in resistance and other important functions, nearly half of the genes in the network correlated specifically with Andigena had unknown gene functions, implying more attention should be drawn to the molecular studies in the primitive potato species in the future.

## Methods

### Raw RNA-Seq Dataset and Normalization of the Gene Expression

The RNA-Seq datasets used for the analysis including differential gene expression and the construction of the network were downloaded from NCBI SRA database Gong et al. (2015); Campbell et al. (2014); Petek et al. (2014); Liu et al. (2015b,a). These RNA-Seq datasets contained 16 tissues of 11 potato varieties and the SRA accessions are shown in Table 1. To simplify the further comparison, only the tissues under the normal growth conditions were selected. The reads of these datasets were filtered using Cutadapt v1.9Martin (2011) and mapped to the potato reference genomeConsortium et al. (2011) (PGSC_DM_v4.03_pseudomolecules.fasta & PGSC_DM_v4.03_genes.gff) from the potato genome database (http://solanaceae.plantbiology.msu.edu/pgsc_download.shtml) using GMAP/GSNAP (http://research-pub.gene.com/gmap/)Wu and Watanabe (2005); Martin (2011). Samtools kit was used to convert the data format and Cufflinks was used to normalize the gene expression levels as fragments per kilobase of exon model per million mapped reads (FPKM)Li (2011); Trapnell et al. (2012). The genes with extremely low expression (FPKM < 3) in average of all tissues were also filtered out in the construction of the co-expression gene network. Differential expression of transcripts was determined using CuffDiff v2.21 Trapnell et al. (2012) and Spearman correlation coefficient analyses between each two datasets were conducted using R packages.

### Gene Co-Expression Network Construction

The gene expression matrix of potato was used to generate the gene co-expression network using weighted gene coexpression network analysis method (WGCNA)Langfelder and Horvath (2008). WGCNA is a systems biology method for describing the correlation patterns among genes across multiple samples and is based on the WGCNA R software package, a comprehensive collection of R functions for performing various aspects of weighted correlation network analysis. The thresholding power *β* was used to calculate adjacency between each two genes and 10 were chosen as optimal thresholding power among a set of candidate powers from 1 to 20 based on the function pickSoftThreshold returned results. The adjacency was calculated and then transformed into Topological Overlap Matrix (TOM) using function TOM similarity and the corresponding dissimilarity was calculated. The gene hierarchical clustering tree was produced using hierarchical clustering setting the minimum module size as 30 and the tree height cut as 0.25. The Module Eigengenes (MEs) were calculated and used to the further analyses. To visualize the network, the edge file and node file of the module were generated for dissimilarity and displayed on Cytoscape softwareShannon et al. (2003).

### Functional Enrichment and Clustering

The Goatools v1.9 based on Python was introduced to analyze the enrichment of genes belonging to the different expression module in order to learn the gene GO terms and functions for each moduleHaibao et al. (2015). The genes participated in network construction were selected to get Gene ontology categories from GO database through Blast2Go (http://www.blast2go.com/b2glaunch/start-blast2go), with parameters of 20 hits and an e-value of 10e^−6^. GO terms conforming to p-value through Bonferroni Correction 0.05 were defined as significantly enriched GO terms. The genome annotation file described above was used as the reference. Only GO terms for Biological Process are shown.

## Acknowledgements

The authors would like to thank Dr.Xianjun Lai (from SCU) for his assistance with some scripts and codes in bioinformatic work. Thanks also to Xia Zhan for scientific English language editorial assistance.

## Author contributions statement

L.Y. and YZ.Z. conceived the analyses, L.Y collected the data, L.Y., Y.Q. and ZR.F. performed the calculations and analysis, L.Y. completed the manuscript. All authors discussed the results and reviewed the manuscript.

